# Inverse correlation between heme synthesis and the Warburg effect in cancer cells

**DOI:** 10.1101/753079

**Authors:** Yuta Sugiyama, Erika Takahashi, Kiwamu Takahashi, Motowo Nakajima, Tohru Tanaka, Shun-ichiro Ogura

**Affiliations:** School of Life Science and Technology, Tokyo Institute of Technology, Yokohama, Kanagawa, Japan; SBI Pharma CO., LTD., Roppongi, Tokyo, 106-6020, Japan

## Abstract

Cancer cells show a bias toward the glycolytic system over the conventional mitochondrial electron transfer system for obtaining energy. This biased metabolic adaptation is called the Warburg effect. Cancer cells also exhibit a characteristic metabolism, a decreased heme synthesizing ability. Here we show that heme synthesis and the Warburg effect are inversely correlated. We used human gastric cancer cell lines to investigate glycolytic metabolism and electron transfer system toward promotion/inhibition of heme synthesis. Under hypoxic conditions, heme synthesis was suppressed and the glycolytic system was enhanced. Addition of a heme precursor for the promotion of heme synthesis led to an enhanced electron transfer system and inhibited the glycolytic system and vice versa. Enhanced heme synthesis leads to suppression of cancer cell proliferation by increasing intracellular reactive oxygen species levels. Collectively, the promotion of heme synthesis in cancer cells eliminated the Warburg effect by shifting energy metabolism from glycolysis to oxidative phosphorylation.

## Introduction

Cancer is caused by accumulation of various genetic mutations. A common feature of most cancer cells is suppression of mitochondrial aerobic respiration and an enhanced glycolytic ATP synthesis for supporting abnormal cellular proliferation and metastasis. Therefore, cancer cells require and utilize abundant glucose and produce excessive lactic acid by accelerated glycolysis, resulting in lactic acidosis [1]. This concept, first advocated by Nobel laureate Otto Warburg in 1924, is widely recognized as the Warburg effect and has pioneered research toward analysis of tumor metabolism.

HIF-1 (hypoxia inducible factor 1) is activated owing to low oxygen concentration in cancer cells within the tumor tissue [2–4]. HIF-1 induces pyruvate dehydrogenase kinase-1 (PDK-1), which inactivates pyruvate dehydrogenase (PDH) [5]. PDH is a key player that converts pyruvic acid to acetyl CoA, which links the glycolytic pathway and TCA cycle. Thus, HIF-1 regulates PDK-1 and functions as a modulator of glycolysis and TCA cycle. Moreover, HIF-1 induces glucose transporter-1 and glycolytic enzymes [6,7] and consequently leads to increased lactic acid formation. Thus, HIF-1 shifts metabolism from oxidative phosphorylation to glycolysis under hypoxic conditions [8,9], thereby promoting the Warburg effect under hypoxia.

5-Aminolevulinic acid (ALA) is a precursor in the porphyrin synthesis pathway leading to heme formation [10]. The rate-limiting enzyme in heme synthesis is the mitochondrial enzyme ALA synthase (ALAS) [11]. ALA is metabolized to protoporphyrin IX (PpIX) by multiple enzymatic reactions while between cytoplasm and mitochondria, and is finally synthesized into heme by the enzyme ferrochelatase, which coordinates iron ion to porphyrin [12]. Administration of ALA into tumors leads to intracellular accumulation of PpIX because cancer cells exert limited ferrochelatase activity [13,14]. Hypoxia is believed to induce a reduction in PpIX accumulation [15,16]. To the best of our knowledge, the influence of hypoxia on heme synthesis has not yet been clarified [17,18].

One of the advantages of the Warburg effect is the ability to suppress cytotoxic reactive oxygen species (ROS), which are produced during aerobic respiration and cause cytotoxicity. Resultantly, cancer cells create a situation favorable for their survival by reducing oxidative stress via suppressing excessive ROS. Recently, attempts have been made to treat cancer by targeting active energy metabolism biased toward cancer glycolysis; for example, studies using dichloroacetic acid (DCA) were reported to induce apoptosis by shifting glucose metabolism from glycolysis to oxidative phosphorylation [19–21].

Our previous research using mouse liver showed that ALA increases activity of the mitochondrial enzyme cytochrome *c* oxidase (COX), a rate-limiting enzyme in the electron transport system (ETS) [22]. Owing to the fact that COX is a heme protein, ALA-mediated increase in the amount of heme was thought to induce the expression COX. We also reported that addition of ALA leads to activation of ETS in cancer cells [23].

Here we investigated metabolic pathways in cancer cells to understand the relationship between heme synthesis and the Warburg effect. To this end, we comprehensively examined the effect of heme synthesis regulation on energy metabolism in cancer cell lines.

## Methods

### Reagents

The substrates 5-aminolevulinic acid (ALA) hydrochloride and succinyl ferrous citrate (SFC) were purchased from Cosmo Bio Co., Ltd. (Tokyo, Japan); 4,6-Dioxoheptanoic acid (Succinylacetone, SA) was purchased from Sigma-Aldrich (St. Louis, MO); and 2,7-Dichlorodihydrofluorescein diacetate (DCFH-DA) and 4,5-dihydroxybenzene-1,3-disulfonate (Tiron) were purchased from Wako Pure Chemical Industries, Ltd. (Osaka, Japan). RPMI-1640 medium and Antibiotic-Antimycotic solution (ABAM, Penicillin-Streptomycin-Amphotericin B mixture) were obtained from Nacalai Tesque (Kyoto, Japan). Fetal bovine serum (FBS) was purchased from Biowest (Nuaillé, France).

### Cell culture

Human gastric cancer cell lines, KatoIII and MKN45, were purchased from RIKEN Bioresource Center (Tsukuba, Japan). Cells were grown in RPMI-1640 medium supplemented with 10% (v/v) heat-inactivated FBS and ABAM and were incubated at 37°C in an incubator with a controlled humidified atmosphere containing 5% CO_2_. Cell culture under hypoxic conditions was performed using AnaeroPack-Kenki 5% (Mitsubishi Gas Chemical Co., Tokyo, Japan).

### Western blot analysis

Western blot analysis was performed as previously described [24]. The following primary antibodies were used: HIF-1α (sc-10790, 1:400 dilution, rabbit polyclonal, Santa Cruz Biotechnology, Dallas, Texas, USA); GLUT1 (ab652, 1:500 dilution, rabbit polyclonal, Abcam, Cambridge, Great Britain); COX IV-1 (sc-58348, 1:200 dilution, mouse monoclonal, Santa Cruz Biotechnology, Dallas, Texas, USA); and Actin (08691001, 1:200 dilution, mouse monoclonal, MP Biomedicals, Santa Ana, United States). The following secondary antibodies were used: horseradish peroxidase (HRP)-conjugated anti-mouse IgG antibody and HRP-conjugated anti-rabbit IgG antibody (1:3000 dilution, Cell Signaling Technology, Beverly, Massachusetts, USA). HRP-dependent luminescence was developed with Western Lightning Chemiluminescent Reagent Plus (PerkinElmer Life and Analytical Sciences, Inc., Waltham, MA, USA) and detected with a Lumino Imaging Analyzer ImageQuant LAS 4000 mini (GE Healthcare UK, Amersham Place, England).

### Measurement of glucose uptake

Fluorescent-labeled glucose, 2-NBDG (2-deoxy-2-[(7-nitro-2,1,3-benzoxadiazol-4-yl)]-D-glucose), was used to measure glucose uptake. Cells (3∼8 × 10^5^ cells/well) were seeded in 6-well plates and preincubated overnight in RPMI-1640 medium in 5% CO_2_ gas at 37°C. After changing the culture medium, cells were incubated under each testing condition for 24 hours. Subsequently, all culture medium was removed and replaced with glucose-free culture medium in the presence or absence of fluorescent 2-NBDG (100 μM). Cells were incubated at 37°C with 5% CO_2_ for 25 minutes and washed three times with PBS. Cells were lyzed with 0.1 M NaOH and extracted by adding an identical volume of perchloric acid: methanol (1:1, v/v) solution. The obtained mixture was centrifuged to pellet proteins (10,000 × *g*, 4°C, 10 minutes). The concentration of 2-NBDG in the supernatant was quantitatively determined by measuring fluorescence in a fluorescence spectrophotometer RF-5300 PC (Shimadzu Corporation, Kyoto, Japan). Excitation and emission wavelengths of 465 nm and 550 nm, respectively, were used to measure fluorescence intensity of 2-NBDG. Glucose uptake levels were normalized by intracellular protein levels measured by Bradford protein assay.

### Measurement of lactate production

Cells were seeded in 96-well plates (2 × 10^4^ cells/well) and preincubated for 2 days in RPMI-1640 medium in 5% CO_2_ gas at 37°C. After changing the culture medium to that devoid of FBS, cells were incubated for 24 hours under each testing condition. Lactate production level was measured using a Glycolysis Cell-Based Assay Kit (600450, Cayman Chemical, Ann Arbor, Michigan, USA) according to the manufacturer’s protocol.

### Measurement of pH change in culture medium

Cells were seeded in 35 mm dishes (3∼8 × 10^5^ cells/well) and preincubated in RPMI-1640 medium overnight in 5% CO_2_ gas at 37°C. After changing to non-buffering medium containing RPMI-1640 powder medium (Sigma-Aldrich) without sodium bicarbonate, cells were incubated for 24 hours under each testing condition. The pH of the culture medium was measured using a pH meter (LAQUAtwin AS-712, HORIBA, Kyoto, Japan) and the H^+^ concentration was calculated from the pH value. The pH of culture medium without cells was measured as background and the value obtained by subtracting the H^+^ concentration of background from the H^+^ concentration of each sample was defined as ΔH^+^. The ΔH^+^ value was normalized to the intracellular protein concentration.

### HPLC analysis of porphyrins

Cells were seeded in 6-well plates and preincubated overnight in culture medium. After the culture medium was changed, cells were incubated for 24 hours under each testing condition. They were washed twice with PBS and treated with 0.1 M NaOH. Cellular lysates were prepared by adding 3-fold volumes of dimethyl formamide (DMF)/2-propanol (100:1, v/v) to oxidize coproporphyrinogen III (CPgenIII) to coproporphyrin III (CPIII). CPIII was detected by HPLC because CPgenIII was unstable and easily oxidized. These mixtures were centrifuged to remove proteins, and the supernatants were incubated for a day at room temperature in the dark. HPLC analysis was performed as previously described [25] with some modifications. Briefly, PpIX and CPIII were separated using an HPLC system (Prominence, Shimadzu, Kyoto, Japan) equipped with a reversed-phase C18 Column (CAPCELL PAK, C18, SG300, 5 μm, 4.6 mm × 250 mm, Shiseido Co., Ltd., Tokyo, Japan). Elution was started with 100% solvent A and 0% solvent B for 5 minutes, with a linear gradient of solvent B (0%–100%) for 25 minutes and later with solvent B for 10 minutes. Flow was maintained at a constant rate of 1.0 mL/min. Porphyrins were continuously detected using a fluorospectrometer (excitation 404 nm; emission 624 nm). Porphyrin concentration was estimated using calibration curves constructed using porphyrin standards.

### HPLC analysis of intracellular heme level

Cells were seeded in 35-mm dishes and preincubated overnight in culture medium. After the culture medium was changed, they were incubated for 24 hours under each testing condition. Cells were washed three times with PBS and treated with 0.1 M NaOH on ice. Elution solvent A contained 1 M ammonium acetate and 12.5% acetonitrile (pH 5.2), and solvent B contained 50 mM ammonium acetate and 80% acetonitrile (pH 5.2). Cellular lysates were prepared by adding 3-fold volumes of elution solvent A/solvent B (1:9, v/v) to extract intracellular heme. These mixtures were centrifuged to remove proteins (10,000 × *g*, 4°C, 10 minutes), and the supernatants were collected. Heme was measured using an HPLC system (Prominence, Shimadzu, Kyoto, Japan) equipped with a reversed-phase C18 Column (CAPCELL PAK, C18, SG300, 5 μm, 4.6 mm × 250 mm, Shiseido Co., Ltd., Tokyo, Japan). Elution was maintained with 10% solvent A and 90% solvent B for 7 minutes. Flow was maintained at a constant rate of 2 mL/min. Heme was continuously detected by absorbance at 404 nm using a spectroscopic detector. Heme concentrations were estimated using calibration curves constructed using heme standard.

### Measurement of COX activity

Measurement of cytochrome *c* oxidase (COX) activity was performed as previously described with some modifications [22]. Mitochondrial fractions were obtained using a Mitochondria Isolation Kit (MITOISO2, Sigma-Aldrich). Briefly, 5 × 10^7^ cells were washed and homogenized in extraction reagent. The homogenate was centrifuged at 600 × *g* for 5 minutes and the supernatant was centrifuged further at 11,000 × *g* for 10 minutes. The pellet was suspended in storage buffer and used as the mitochondrial fraction. Protein concentrations were determined by the Bradford assay (Bio-Rad Laboratories, CA). COX activity was measured using a Cytochrome *c* Oxidase Assay Kit (Sigma-Aldrich). Briefly, 100 μg of the mitochondrial fraction was diluted with enzyme dilution buffer containing 1 mM n-dodecyl β-d-maltoside. Ferrocytochrome *c* (reduced cytochrome *c* with dithiothreitol) was added to the sample, and COX activity was measured by the decrease in absorption at 550 nm. The difference in extinction coefficients between reduced and oxidized cytochrome *c* is 21.84 at 550 nm [26]. One unit of COX activity was defined as the oxidization of 1.0 μmole of ferrocytochrome *c* per minute at pH 7.0 at 25°C.

### Determination of mitochondrial DNA copy number

Genomic DNA was isolated using the NucleoSpin^®^ RNA II and NucleoSpin^®^ RNA/DNA Buffer Set kit (MACHEREY-NAGEL, Düren, Mannheim, Germany) kit according to the manufacturer’s instructions. Quantitative real-time PCR was performed with SYBR Premix Ex Taq (TaKaRa, Shiga, Japan) using Thermal Cycler Dice^®^ Real Time System Single (TaKaRa, Shiga, Japan) to determinate the amount of mitochondrial DNA (mtDNA) content relative to the nuclear DNA (nuDNA). Primer sets were as follows; human mtDNA, forward 5’-CACCCAAGAACAGGGTTTGT-3’ and reverse 5’-TGGCCATGGGTATGTTGTTA-3’; human nuDNA, forward 5’-TGCTGTCTCCATGTTTGATGTATCT-3’ and reverse 5’-TCTCTGCTCCCCACCTCTAAGT-3’. The PCR amplification conditions included 95°C for 30 seconds; 50 cycles at 95°C for 5 seconds and 60°C for 60 seconds each; dissociation for at 95°C for 15 seconds and at 60°C for 30 seconds; and at 95°C for 15 seconds on a Thermal Cycler Dice Real-Time System. Thermal Cycler Dice Real-Time System analysis software (TaKaRa, Shiga, Japan) was used for data analysis. The Ct values (cycle threshold) were calculated using the crossing-point method, and the relative mtDNA and nuDNA levels were measured by comparison with a standard curve. The mtDNA content was normalized with the content of nuDNA.

### Cell growth assay

MKN45 cells (8 × 10^4^ cells) were incubated overnight at 37°C in 5% CO_2_ in RPMI-1640 medium. The culture medium was changed to fresh medium containing 1 mM ALA, 0.5 mM SFC, and/or 100 µM Tiron and cultured for up to 4 days in 5% CO_2_ in dark. Cells were collected after trypsin treatment, and the number of living cells was determined using trypan blue dye exclusion assay.

### Detection of reactive oxygen species

ROS detection assay was performed using the cell-permeable fluorogenic probe DCFH-DA. Briefly, DCFH-DA diffuses into cells and is deacetylated by cellular esterase to DCFH, which is rapidly oxidized to highly fluorescent DCF by ROS. The fluorescence intensity of DCF can be assessed as an indicator of cellular ROS levels.

MKN45 cells were incubated with 1 mM ALA, 0.5 mM SFC, and/or 100 µM Tiron for 24 hours. After washing with PBS, the media was changed to serum-free medium supplemented with 10 µM DCFH-DA. After incubating for 30 minutes, the medium was discarded and cells were collected by a cell scraper using 200 µL of Hanks’ balanced salt solution (HBSS) (-). The collected solution was suitably diluted with HBSS (-) and the fluorescence was measured using a spectrophotometer F-7000 (Hitachi High-Tech Science, Tokyo). DCF fluorescence intensity was detected at an excitation wavelength of 480 nm and a fluorescence wavelength of 525 nm.

### Statistical analysis

Data are expressed as means ± standard deviation using two or three independent experiments. Data were analyzed for statistical significance using Student’s t-test for the comparison between the control and experimental groups. Difference was assessed with two-sided test with an α level of 0.05. In case of three or more groups, statistical significance was analyzed using the Dunnett’s test or Tukey-Kramer’s test with an α level of 0.05. Statistical analyses were performed using the JMP^®^ 13 software (SAS Institute Inc., Cary, NC, USA).

## Results

### Glycolytic increase in gastric cancer cells under hypoxia

We examined the gastric cancer cell lines KatoIII and MKN45 to understand the relation between glycolysis and heme synthesis under hypoxia. Under hypoxic conditions, cancer cells induce expression of the glucose transporter GLUT1 and increase glucose uptake via HIF-1, thereby resulting in excessive lactic acid production and shifting the acid-base equilibrium toward the acidic side [6]. Therefore, we assessed the functional status of the glycolytic system by measuring the expression of GLUT1, uptake of a fluorescent glucose analog, 2-NBDG, lactic acid concentration, and pH of the culture media.

The cell lines were cultured for 24 hours under normoxic (21% O_2_) or hypoxic (1% O_2_) conditions. On performing expression analysis, in both cell lines, the expression of GLUT1 was found to be enhanced by HIF-1α under hypoxic conditions (Fig 1A). In correlation with up-regulated GLUT1, the uptake of 2-NBDG was significantly increased under hypoxic conditions (Fig 1B). Moreover, concentrations of lactic acid concentration (Fig 1C) and H^+^ concentration (Fig 1D) were also significantly increased. Thus, hypoxic conditions accelerated glycolysis in gastric cancer cell lines. These experiments also served to confirm their application in evaluating the glycolytic system.

**Fig 1.**
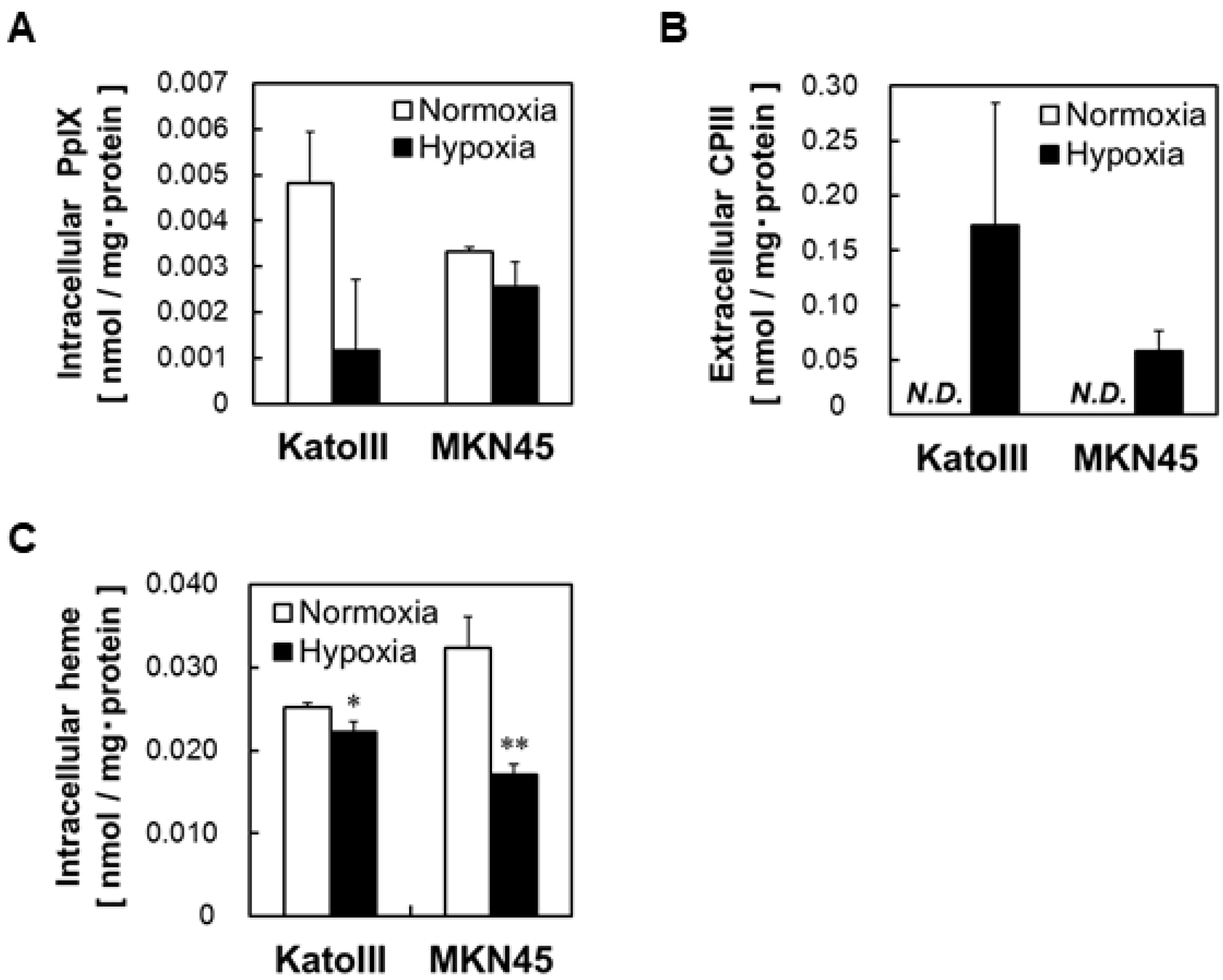
Hypoxia enhances glycolysis in human gastric cancer cell lines. KatoIII and MKN45 cells were used for experiments following incubation either under normoxia (21% O_2_) or hypoxia (1% O_2_) for 24 hours. (A) HIF-1α and GLUT1 were detected by Western blotting. (B) Glucose uptake was assessed after incubation for 25 minutes with 100 μM 2-NBDG. (C) Extracellular lactate concentration and (D) change in pH of medium was measured. Values are means ± SD, n = 3, Asterisk indicates a significant difference (*; *p* < 0.05, **; *p* < 0.01).

### Reduction of intracellular PpIX and heme levels in gastric cancer cells under hypoxic conditions

Previous studies have shown that under hypoxic conditions, the accumulation of PpIX reduces when ALA is supplemented [15,16]. However, the effect of hypoxia on porphyrin synthesis or accumulation, without ALA supplementation, has not yet been clarified. Therefore, we measured the amount of the porphyrin in gastric cancer cell lines by high-performance liquid chromatography (HPLC) after culturing for 24 hours under normoxic (21% O_2_) or hypoxic (1% O_2_) conditions. The amount of intracellular PpIX in both KatoIII and MKN45 cells was reduced under hypoxic conditions (Fig 2A), whereas extracellular coproporphyrin III (CPIII) was remarkably increased (Fig 2B). Conversely, no other type of porphyrin was detected intra- or extra-cellularly. Thus, under hypoxic conditions, the porphyrin synthesis pathway is thought to be inhibited at the CPgenIII step resulting in excretion of the synthesized CPgenIII and decrease in PpIX. The amount of intracellular heme under hypoxia was also measured by HPLC after culture for 24 hours under normoxic (21% O_2_) or hypoxic (1% O_2_) conditions (Fig 2C). A significantly reduced intracellular heme concentration indicated inhibitory effect of hypoxia on heme synthesis. This also supported the hypothesis that the porphyrin synthesis pathway is inhibited at the CPgenIII step under hypoxia. Thus, we can postulate a negative correlation between heme synthesis and the glycolytic system.

**Fig 2.**
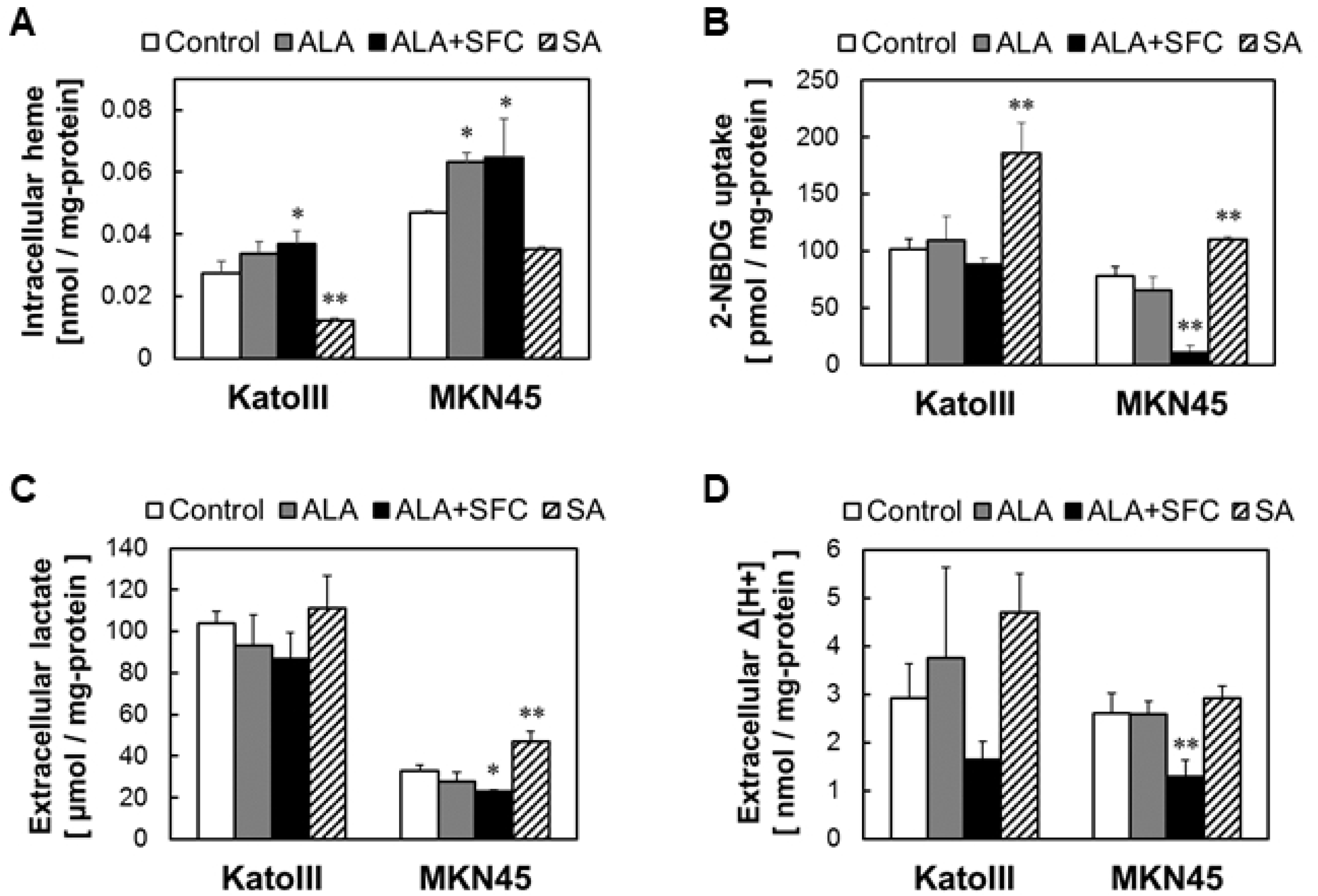
Hypoxia leads to excretion of the heme-intermediate CPIII and reduction of heme content in human gastric cancer cell lines. KatoIII and MKN45 cells were incubated under normoxia (21% O_2_) or hypoxia (1% O_2_) for 24 hours and intracellular PpIX (A), extracellular CPIII (B) and intracellular heme concentration (C) were measured. Other porphyrins were not detected. Values are means ± SD, n = 2 (A, B) and n = 3 (C), Asterisk indicates a significant difference (*; *p* < 0.05, **; *p* < 0.01).

### Inverse correlation between heme biosynthesis and glycolysis

Next, we examined the effect of heme synthesis on glycolysis in cancer cell lines using ALA, SFC as iron ion, and SA. The addition of ALA and iron ion promotes heme synthesis because the rate-limiting step in PpIX production is ALA synthesis and heme formation by coordination of divalent iron ions to PpIX [11]. Conversely, SA suppresses heme synthesis by inhibiting ALA dehydratase, which functions in condensing two molecules of ALA.

The gastric cancer cell lines were cultured for 24 hours in media containing 1 mM ALA, 0.5 mM SFC, or 0.5 mM SA, followed by measurement of intracellular heme by HPLC. The addition of ALA alone or ALA+SFC induced an increase in intracellular heme concentration, whereas SA led to its reduction (Fig 3A). Thus, heme synthesis can be regulated by the addition of ALA, SFC, or SA.

**Fig 3.**
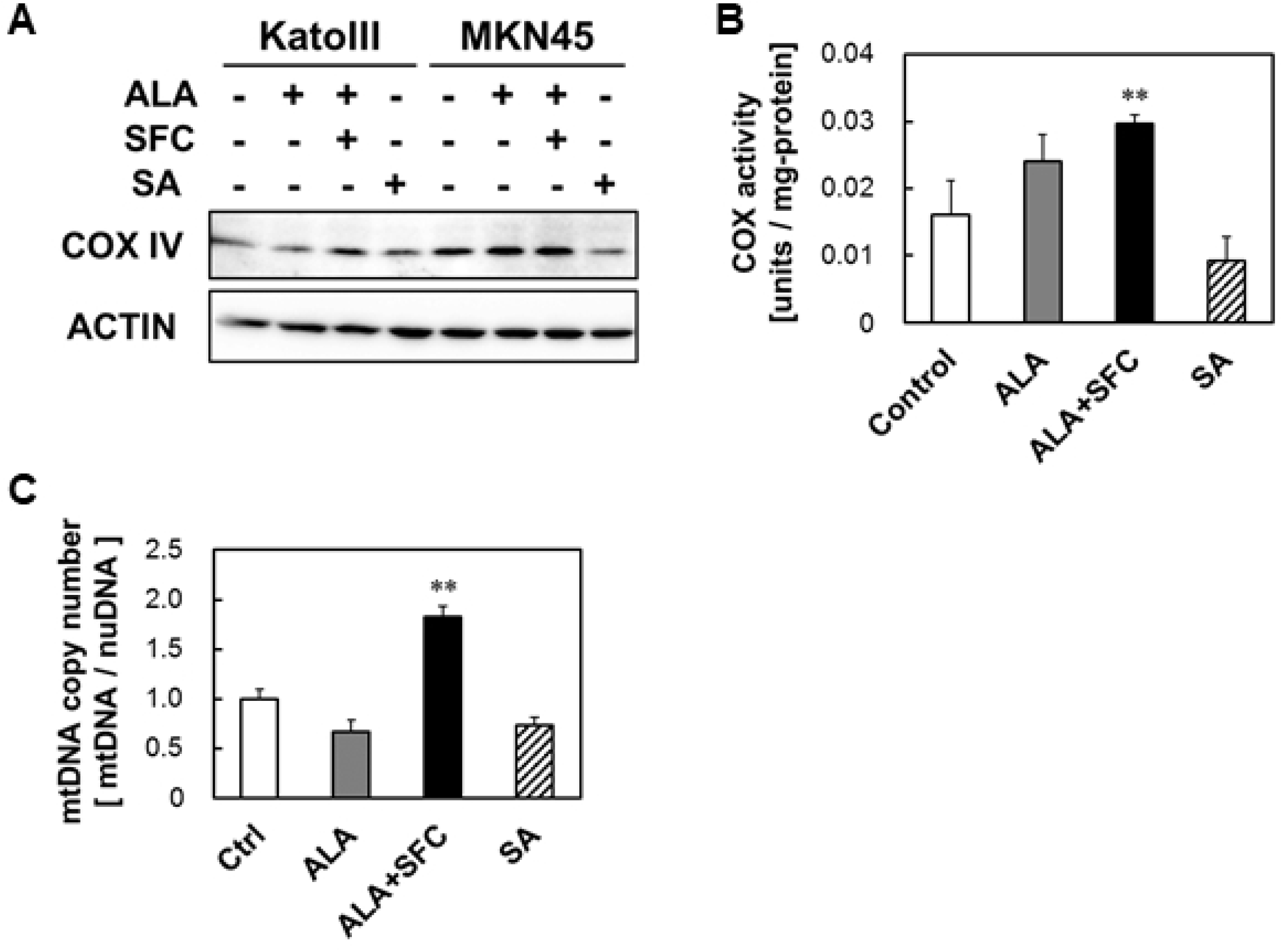
Heme synthesis is inversely correlated with glycolysis. KatoIII and MKN45 cells were used for experiments following incubation with 1 mM ALA, 0.5 mM SFC and 0.5 mM SA for 24 hours. (A) Intracellular heme concentration measured by HPLC. (B) Glucose uptake assessed after 25 minutes of incubation with 100 μM 2-NBDG. (C) Extracellular lactate concentration and (D) change in pH of medium was measured. Values are means ± SD, n = 3, Asterisk indicates a significant difference analyzed by Dunnett’s test (*; *p* < 0.05, **; *p* < 0.01).

Next, we tested the effect of heme synthesis on glycolysis by analyzing the uptake of 2-NBDG, lactic acid concentration, and H^+^ concentration after 24 hours of cell culture with 1 mM ALA, 0.5 mM SFC, or 0.5 mM SA. Under conditions that enhanced heme synthesis (i.e., supplementation with ALA or ALA+SFC), the uptake of 2-NBDG was similar or reduced compared with control (Fig 3B). Conversely, the uptake of 2-NBDG was significantly higher under conditions that suppressed heme synthesis (SA). Lactic acid concentration in MKN45 cells was down-regulated by supplementation of ALA or ALA+SFC and up-regulated by addition of SA (Fig 3C). However, no such difference was observed in KatoIII cells. Furthermore, both cell lines showed a tendency toward decrease in H^+^ concentration upon addition of ALA+SFC (Fig 3D). H^+^ concentration in KatoIII cells was increased after supplementation with SA (Fig 3D). Thus, heme synthesis and glycolysis are inversely related in gastric cancer cell lines.

### Correlation between heme synthesis and mitochondrial activity

COX, a heme protein, is the rate-limiting enzyme of the mitochondrial electron transfer chain [27,28]. Injection of ALA induced expression and activity of COX in mice liver [22]. It also enhanced COX expression and oxygen consumption in a lung cancer cell line, A549 [23]. Accordingly, we expected that regulation of heme synthesis using ALA, SFC, and SA should affect mitochondrial activity.

First, we examined the expression of COX to investigate the effect of heme synthesis regulation on mitochondrial activity. We evaluated the expression of COX IV-1, the nuclear-encoded largest subunit of COX and isoform expressed under normal conditions [29], in cancer cells after incubation for 24 hours with ALA, SFC, and SA (Fig 4A). COX expression increased in both cell lines when heme synthesis was enhanced by ALA or ALA+SFC. In contrast, COX expression was reduced in MKN45 cells when heme synthesis was suppressed by SA. Analysis of COX activity in MKN45 cells revealed significant elevation upon ALA and ALA+SFC supplementation and reduction after SA supplementation (Fig 4B). These results indicate that the expression (Fig 4A) and enzyme activity of COX increased when heme synthesis was promoted and decreased when it was suppressed. Enhanced heme synthesis after addition of ALA+SFC led to up-regulation of COX, most probably because it is a hemoprotein. It can thus be predicted that SA-mediated suppression of heme synthesis would probably reduce COX expression and activity.

**Fig 4.**
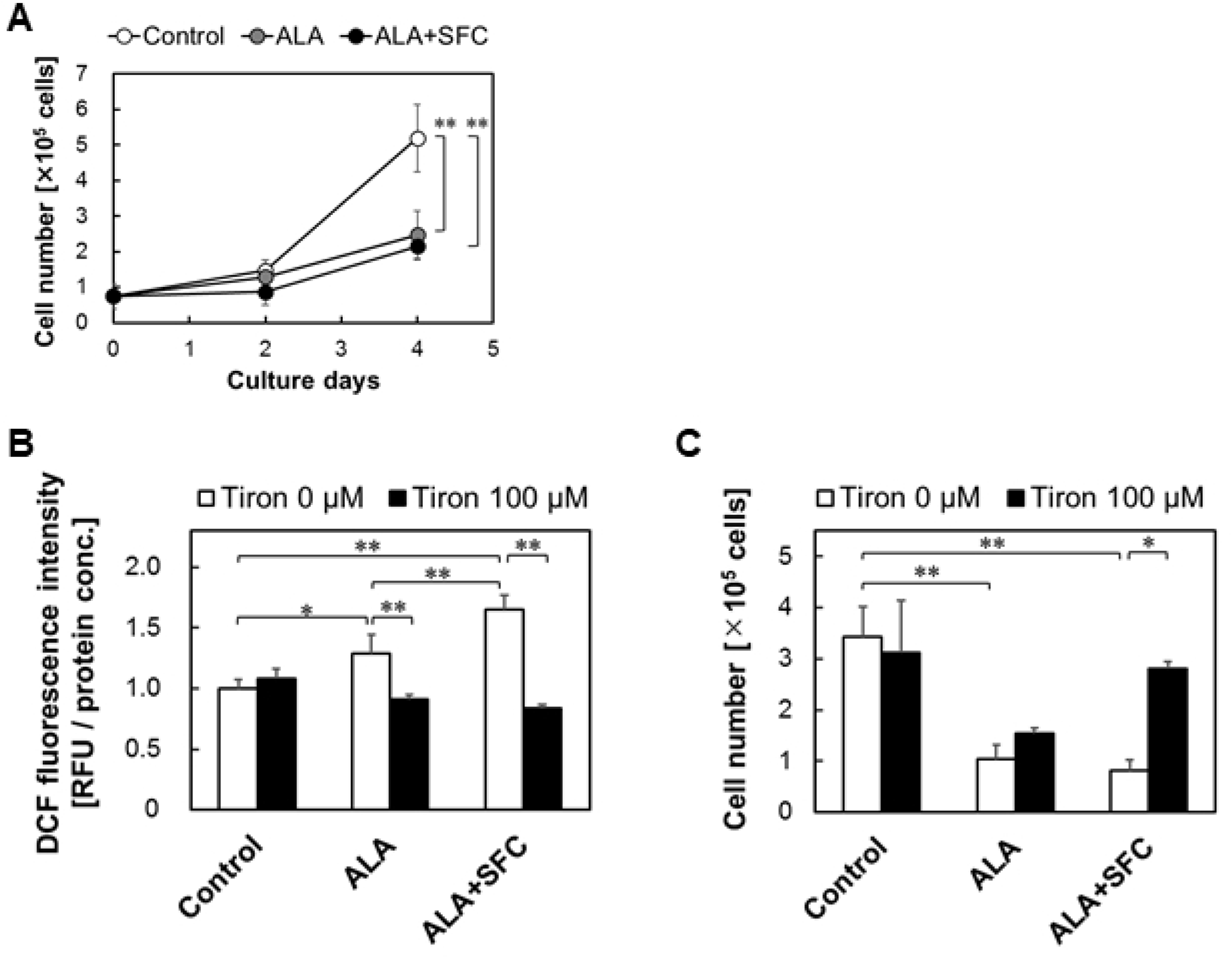
Heme synthesis is positively correlated with COX expression and mtDNA copy number. (A) Expression of COX IV in KatoIII and MKN45 cells was detected by Western blotting after incubation with 1 mM ALA, 0.5 mM SFC, and 0.5 mM SA for 24 hours. (B) MKN45 cells were incubated with 1 mM ALA, 0.5 mM SFC, and 0.5 mM SA for 24 hours; the cells were then harvested and COX activity was measured. (C) Relative copy number of mitochondrial DNA (mtDNA) versus nuclear DNA (nuDNA) of MKN45 cells was determined by real-time PCR. Cells were treated with 1 mM ALA, 0.5 mM SFC, and 0.5 mM SA for 24 hours before the test. The relative mtDNA copy number of the control is shown as 1 in the graph. Values are means ± SD, n = 3 (B) and n = 2 (C), Asterisk indicates a significant difference analyzed by Dunnett’s test (*; *p* < 0.05, **; *p* < 0.01).

Second, we measured the copy number of mitochondrial DNA (mtDNA) (Fig 4C). Genomic DNA was extracted from MKN45 cells after cell culture for 24 hours under each testing condition (1 mM ALA, 0.5 mM SFC, and 0.5 mM SA). Quantitative real-time PCR was performed to determinate the amount of mtDNA relative to the nuclear DNA (nuDNA). The mtDNA copy number increased upon supplementation of ALA+SFC and decreased after addition of SA. Thus, mitochondrial concentration can be correlated with enhanced heme synthesis.

This further indicated a positive correlation between synthesis of heme and activity of mitochondria, including that of COX. Thus, enhanced heme synthesis is suggested to resolve the Warburg effect by increasing COX activity and vice versa.

### Inhibition of cancer proliferation via ROS by promoting heme synthesis

Next, we examined the proliferation of cancer cells under conditions where heme synthesis was enhanced and the Warburg effect was eliminated. MKN45 cells were cultured with ALA and/or SFC for 4 days, following which the number of cells was counted (Fig 5A). Addition of ALA and ALA+SFC led to suppression of cell proliferation to <50% compared with the control group.

**Fig 5.**
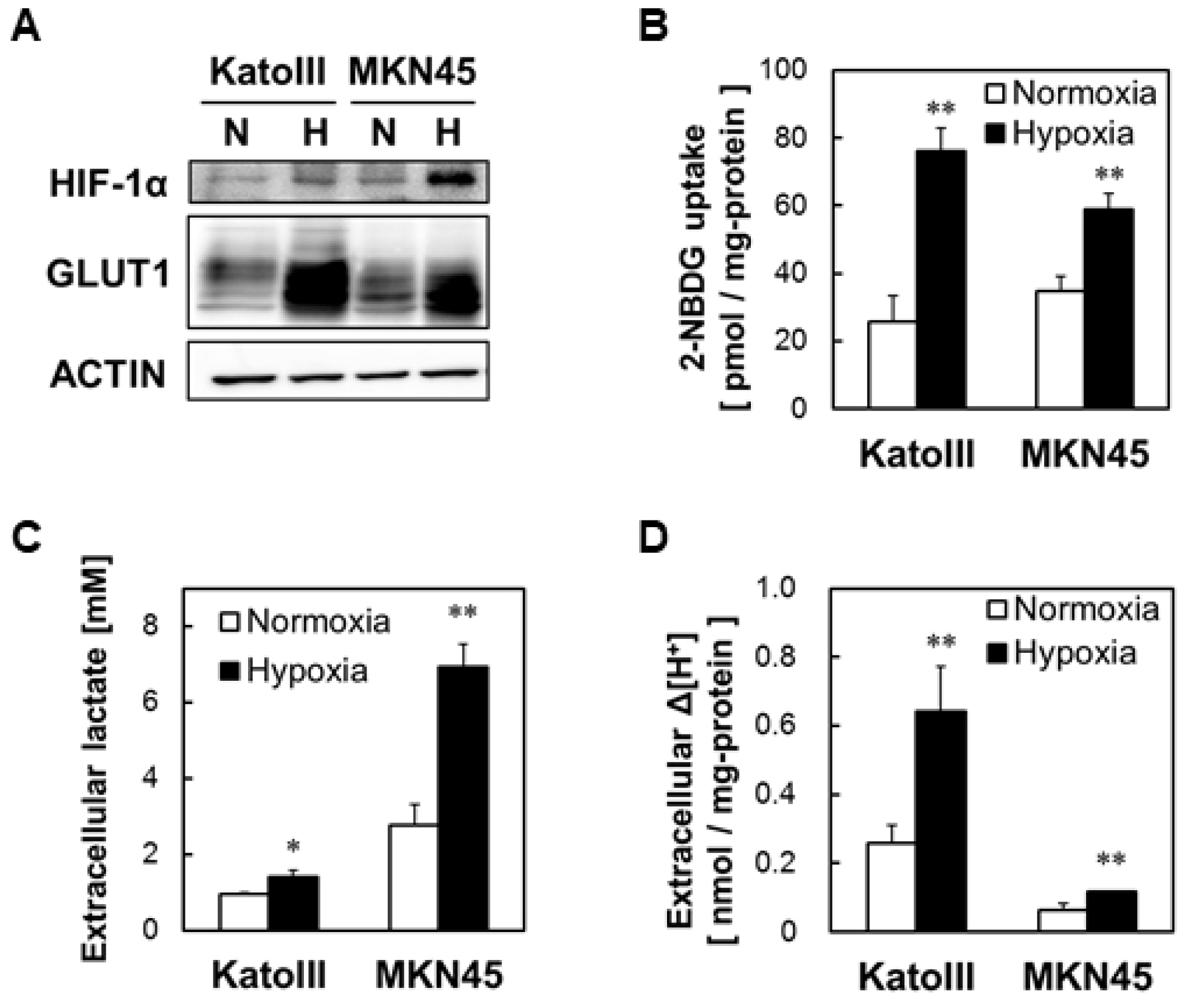
MKN45 proliferation was inhibited by ALA+SFC and it was canceled by addition of a ROS scavenger Tiron. (A) MKN45 cell numbers estimated by trypan blue dye exclusion assay on day 0, 2, 4. (B) MKN45 cells were incubated with 1 mM ALA, 0.5 mM SFC and 100 µM Tiron for 24 hours and intracellular ROS levels were detected using a DCFH-DA based method. The graph shows relative fluorescence intensity of DCF. (C) MKN45 were cultured with 1 mM ALA, 0.5 mM SFC, and 100 µM Tiron for 24 hours and growth assay was performed similar to that in Fig 5a. Values are means ± SD, n = 3, Asterisk indicates a significant difference analyzed by Dunnett’s test (A) and Tukey-Kramer’s test (B, C) (*; *p* < 0.05, **; *p* < 0.01).

Mitochondria, the seat of oxidative phosphorylation, are also the main source of ROS generation [30,31]. The amount of ROS was measured to investigate the increased COX activity upon addition of ALA (Fig 5B). The changes in the amount of ROS were also determined when Tiron was added as a ROS scavenger [32,33]. ROS increased significantly after addition of ALA and/or SFC and was markedly suppressed under conditions where Tiron was added. MKN45 cells were cultured with ALA, SFC, and Tiron for 4 days and the growth was measured (Fig 5C). The ALA+SFC-mediated suppression of cell growth was significantly neutralized by Tiron. Thus, the proliferation of cancer cells was suggested to be suppressed by intracellular ROS caused by ALA+SFC.

Thus, ALA+SFC were suggested to promote heme synthesis, enhance aerobic respiration, generate ROS, and inhibit proliferation of cancer cells.

## Discussion

In this study, we evaluated energy metabolism in cancer cells to better understand the relationship between heme synthesis and the Warburg effect. Indicators of glycolysis namely, GLUT1 expression, uptake of 2-NBDG, lactic acid production, and pH of culture media were significantly up-regulated under hypoxic conditions. Further, heme metabolism and intracellular heme concentration was significantly reduced under hypoxic conditions. Thus, a negative correlation between heme synthesis and glycolysis was suggested based on hypoxia-induced acceleration of the glycolytic system and deceleration of heme synthesis in gastric cancer cell lines. The effect of ALA+SFC and SA, up- and down-regulators of heme metabolism, respectively, on COX and mitochondrial DNA suggested that promotion of heme synthesis led to a shift in energetic metabolism from glycolysis to oxidative phosphorylation by enhancing COX expression and activity of cancer cells. This led to elimination of the Warburg effect. Finally, gastric cancer cells exposed to ALA+SFC showed significantly reduced cell growth caused by increased intracellular ROS generation.

ALA is a naturally occurring amino acid synthesized *in vivo* from succinyl CoA and glycine and is metabolized to porphyrin in multiple steps occurring in the cytoplasm and mitochondria. Ferrochelatase catalyzes the insertion of ferrous iron into PpIX and is thereby the terminal step of heme synthesis and conversion of PpIX to heme. Similarly, tumor cells are known to accumulate PpIX under low activity of ferrochelatase [13,14]. This phenomenon allows photodynamic diagnosis using ALA [34,35]. Earlier, we reported that the membrane transporter ABCB6, which is mainly expressed on mitochondrial membrane for transporting CPgenIII from cytosol into mitochondria, is polarized on the cytoplasmic membrane under hypoxic conditions. This results in the extracellular export of porphyrins [25]. Our results (Fig 2AB) are consistent with this mechanism.

Earlier, we have shown that COX activity and ATP concentration was strongly up-regulated in mouse liver after ALA was injected [22]. In this study, the human gastric cancer cell line MKN45 showed increased expression and activity of COX in presence of ALA+SFC. Also, ALA+SFC act as heme substrates and lead to up-regulation of COX in both normal and cancer cells. This is supported by that the inhibitor of heme synthesis, SA, suppressed COX up-regulation and elevated glycolysis (Fig 3, 4).

Most cancer cells suppress mitochondrial aerobic respiration and synthesize ATP necessary for proliferation and metastasis by enhancing the glycolytic system. Thus, cancer cells require a large amount of glucose. This was advocated by Warburg in 1924 and became widely recognized as the Warburg effect [1]. Glycolysis is not efficient for ATP production but forceful for other substance productivity such as nucleic acids and lipids [36–38]. Therefore, glycolysis is better than oxidative phosphorylation for rapid proliferation. Several attempts have been made to treat cancer cells by focusing on this characteristic metabolic adaptation of cancer cells [39–41]. Glucose derivatives such as fluorodeoxyglucose (FDG) and 2-deoxyglucose (2-DG) are used for diagnosis and treatment of cancer [41–43].

Attempts have been made to treat cancer by inducing ROS production from mitochondrial respiration such as dichloroacetate (DCA). The conversion of pyruvate to acetyl CoA by pyruvate dehydrogenase (PDH) is the rate-limiting step in glycolysis. DCA activates PDH and inhibits lactic acid synthesis and is therefore used as an oral therapeutic agent for mitochondrial diseases, including lactic acidosis. It has reported that DCA restores PDH activity of cancer cells and activates oxidative phosphorylation and consequently induces apoptosis [19,21]. Besides, it is reported that the small molecule ATN-224, which is an inhibitor of superoxide dismutase 1 (SOD1), leads to reduction of lung tumor volume *in vivo* and ROS-dependent cell death *in vitro*. Thus, controlling ROS can be effective in cancer treatment.

It is known that superoxide anion radicals are generated by trapping electrons that leak out of oxygen molecules when the mitochondrial membrane potential becomes unstable owing to changes in the activity of ETS [44,45]. The activation of COX in MKN45 cells by ALA+SFC (Fig 4) indicated that the leakage of superoxide anions from mitochondria most probably led to increased ROS (Fig 5). We used Tiron, an analog of vitamin E and a radical scavenger, as a ROS scavenger [32,33]. Tiron reduced ROS levels and showed recovery of cell proliferation (Fig 5). Taken together, it suggested that ALA+SFC enhanced mitochondrial activity and leakage of superoxide anion radicals followed by an induction in cancer cell death.

To the best of our knowledge, there is no comprehensive study on the relationship between heme synthesis and the Warburg effect. This study provides a novel insight into the complex metabolism of cancer cells and presents a promising new therapeutic strategy for treating cancer. Further studies are required to reveal the molecular mechanism of ALA combined with the Warburg effect toward disruption and elimination of cancer cells.

